# The genomic legacy of selectively breeding rhesus macaques for HIV/AIDS-related research

**DOI:** 10.1101/2025.05.26.654976

**Authors:** MM Lyke, A Bagwell, D Newman, S Galindo, T Church, C Christensen, SB Gray, LA Cox, CN Ross, D Kaushal, IH Cheeseman, SA Cole

## Abstract

**Background:** Rhesus macaques are the preferred non-human primate model for HIV/AIDS-related research. Specific major histocompatibility complex (MHC) haplotypes and genetic ancestry are linked to slow disease progression of Simian Immunodeficiency Virus (SIV) infections in macaques. To maximize their utility for HIV/AIDS research, the Southwest National Primate Research Center (SNPRC) has introduced targeted breeding strategies to reduce the prevalence of SIV-refractory MHC haplotypes and levels of admixture between Indian- and Chinese-origin macaques.

**Results:** We characterized the MHC region in SNPRC macaques by targeted deep sequencing of 1,458 animals born over a ten-year period. Following the implementation of MHC management strategies, the prevalence of SIV-refractory MHC haplotypes reduced significantly while overall haplotype diversity was maintained. We investigated the impact of management strategies on admixture, population genetic structure, genetic diversity and inbreeding using whole exome sequencing of founding and colony-born animals (n=488). Admixture analysis of founders showed some Chinese ancestry, though population substructure more closely reflected primate research center source than geographical origin. The levels of Chinese ancestry declined significantly over time, though genetic diversity remains high. Finally, we performed genome-wide scans for genetic selection over time. We identify numerous genomic regions where allele frequencies have shifted significantly, supporting the presence of short-term adaptation within the colony.

**Conclusions:** We show that colony management strategies have been successful without reducing genetic diversity of the MHC or exonic regions. We also show that colony genetic substructure is related to animal colony source and that mergers and migrations have reduced inbreeding and increased overall genetic diversity.

## Background

Rhesus macaques (*Macaca mulatta*) are the preferred non-human primate (NHP) model for translational research. They are closely related and physiologically similar to humans, making macaques highly sought after models for many biomedical applications, including infectious disease research (1–5). Rhesus macaques have been integral in the investigation of viral, bacterial and parasitological disease, contributing to our understanding of pathogenic transmission, disease progression, and in the development and testing of drugs and vaccines (1–6). Notably, rhesus macaques have long been the predominant animal model for studies of human immunodeficiency virus (HIV) and acquired immunodeficiency syndrome (AIDS), which remains one of the most widespread and pervasive infectious diseases worldwide (1, 5, 7–9). Rhesus macaques challenged with simian immunodeficiency virus (SIV) show similar disease progression and pathogenicity as humans infected with HIV and are thus considered an effective surrogate translational model (1, 7, 8, 10).

Despite their widespread use, special considerations are needed when using rhesus for HIV/AIDS-related research. In rhesus macaques, some major histocompatibility complex (MHC) class I haplotypes are associated with reduced viral load and slow disease progression upon SIV-infection (7, 11–20). Subsequently, rhesus designated for HIV/AIDS-related research are routinely sequenced to identify and exclude these MHC haplotypes. In 2000, the National Institutes of Health (NIH) introduced support for the breeding and maintenance of specific pathogen-free (SPF) Indian-origin rhesus macaques to support ongoing HIV/AIDS research (8). The SPF rhesus macaque colony housed at the Southwest National Primate Research Center (SNPRC) is funded by this mechanism, therefore the genetic management of the colony must incorporate MHC characterization. To meet this requirement, the SNPRC began routinely sequencing the rhesus MHC region through a collaboration with the Wisconsin National Primate Research Center (WNPRC) Genomics Services (https://primate.wisc.edu/research-services/genomics-services/). Four MHC haplotypes that are associated with slowed disease progression (*Mamu-A001*, *Mamu-B003*, *Mamu-B008*, *Mamu-B017*) were present in the colony, leading to management strategies aimed to reduce animals with those haplotypes in the breeding population and to increase production of animals negative for all four haplotypes (referred to as “quad negatives”).

Another important consideration is the ancestry and genomic admixture of rhesus macaques designated for HIV/AIDS research. Rhesus macaques inhabit the broadest geographical range of any NHP species, and, due to historical periods of isolation, local adaptations have led to genetic and morphological divergence (21–24). Genetic evidence indicates that regional biological distinctions in immune response can directly impact their suitability as biomedical research models (23, 24). Historically, rhesus used for research were primarily of Indian origin, though their export was banned in 1978 (25). This led to the expansion of Indian-origin rhesus breeding programs in the United States (US) and China became the largest rhesus exporter (26). Thus, most breeding facilities house either Indian- or Chinese-origin rhesus macaque colonies. Since the founding of US-based rhesus breeding colonies many facilities have experienced unintentional admixture between Indian- and Chinese-origin animals, leading to the development of tools to genetically distinguish between them (27). While admixed animals are suitable for some research initiatives, Indian- and Chinese-origin rhesus respond differently to SIV infection, making non-admixed animals of either origin preferable for HIV/AIDS-related research (28–30).

In addition to ancestral origin, local sources of animals can also impact the genetic substructure of the population (31). Recent demographic history can influence rates of evolution through gene flow, genetic drift and selection, impacting the genomic distribution of genetic variation (32–35). Population substructure can also cause false-positive results or reduce power in disease-related genome-wide association studies (36, 37), making it important to analyze animals designated for biomedical research. There are three primary sources of animals in the SNPRC SPF rhesus macaque colony. The colony was founded with animals transferred from Brooks Air Force Base (BAFB) in the early 2000s, though there was a separate conventional (non-SPF) colony on campus prior to that time. SPF-derived animals from the conventional colony were transferred into the primary colony starting in 2007. However, it was otherwise a closed colony until 2017 when it merged with ≈240 rhesus imported from the New England National Primate Research Center (NEPRC). Previous research has indicated genetic substructure in rhesus macaques housed at various research centers (38), signifying that there may be substructure in the SNPRC colony due to genetic variation between founding animals and sources of immigrations.

The primary goals of this study are to investigate ways in which genetic colony management strategies and source-population genetic dynamics have impacted the genetic composition and suitability of the SNPRC colony for HIV/AIDS-related research. We hypothesized that: 1) by selecting against certain MHC haplotypes in our breeding animals, MHC diversity would decrease and result in the loss of rare alleles; 2) genetic variation between source populations would lead to population substructure; 3) without new migrations, overall genetic diversity would decrease over time in the rhesus colony; 4) by merging new animals into SNPRC breeding groups, the genetic diversity would increase in subsequent generations. Here we combine extended MHC haplotype sequencing, whole exome sequencing, and robust bioinformatic analyses to test these hypotheses and to characterize our rhesus macaque colony for overall genetic and MHC diversity appropriate for HIV/AIDS-related research.

## Results

### Selective breeding has reduced the frequency of SIV refractory MHC haplotypes

We obtained extended MHC haplotypes by targeted deep sequencing of 1,458 rhesus macaques, including full birth cohorts between 2012 and 2022, from the WNPRC Genomics Services (14). We estimated haplotype diversity by birth year across this robust dataset and saw no significant change in overall *Mamu-A* or *Mamu-B* haplotype diversity across the study period (p=0.42; Figure 1B). For the four monitored haplotypes (*Mamu-A001*, *Mamu-B003*, *Mamu-B008*, *Mamu-B017*), we successfully removed all animals with the B003 haplotype (*n* = 7) from breeding in 2017. *Mamu-A001* is one of the four most common *Mamu-A* haplotypes among rhesus and has consequently remained fairly consistent in the population, with a recent sharp decrease in frequency from 17.3% of all haplotypes for animals born in 2021 to 8.3% in 2022 (Fig. 1C). *Mamu-B008* and *Mamu-B017* have both decreased gradually, with *Mamu-B008* decreasing from 4.5% to 1.8% from 2012 to 2022 and *Mamu-B017* decreasing from 14.1% to 6.6% during that same time (Fig 1C). Figure 1D shows the percentage of total births that were negative for *Mamu-A001*, *Mamu-B003*, *Mamu-B008*, *Mamu-B017* from 2012-2022. This shows a significant increase over time (mean increase per year of 1.25%, *p* = 0.042, linear model) from 39% of births in 2012 to 61% in 2022 (Fig 1D).

**Figure 1.**
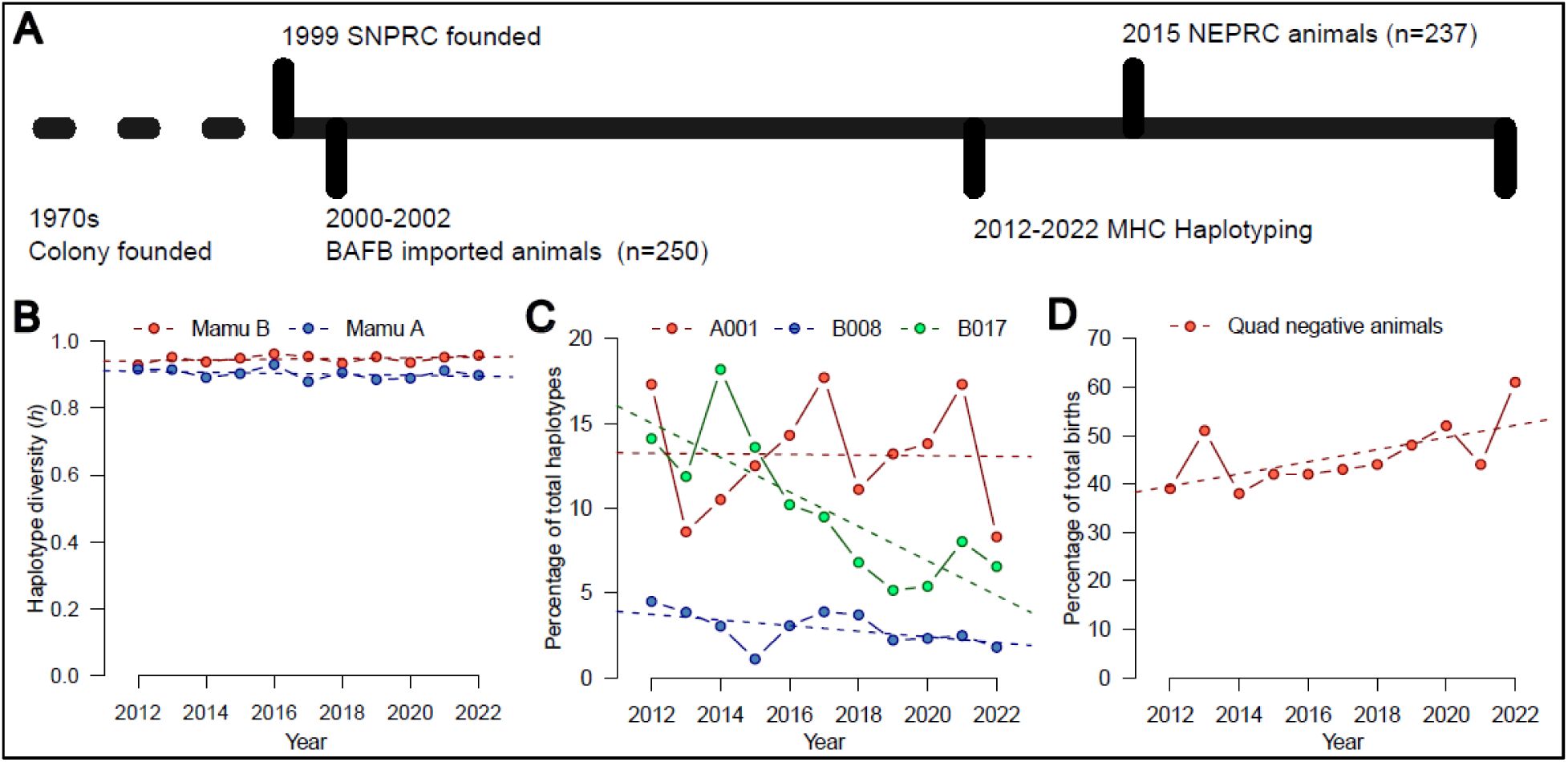
Monitoring MHC haplotypes in the SNPRC rhesus macaques. (A) A timeline of colony mergers and MHC haplotype monitoring. (B) Average *Mamu-A* and *Mamu-B* haplotype diversity by birth year. (C) Percentage of monitored haplotypes *Mamu-A001*, *Mamu-B008*, *Mamu-B017* relative to total haplotypes by birth years from 2012-2022. (D) Percentage of total births negative for all monitored MHC haplotypes (quad negative) from 2012-2022.

### Founding populations show distinct patterns of admixture, diversity and inbreeding

Over the last ten years we have successfully reduced SIV-refractory MHC haplotypes at SNPRC. To understand the impact of this management strategy, we characterized the genomes of animals from founding cohorts using whole exome sequencing (WES). We have previously demonstrated that WES using exome capture kits designed for the human genome performed effectively with coverage of a high proportion of the macaque exome (39). We generated WES data for 212 founding animals from the conventional source (*n* = 28), Brooks source (*n* = 97), and the NEPRC source (*n* = 87). After quality control filtering we retained 917,713 high quality biallelic variants with a mean minor allele frequency of 0.08.

Principal component analysis (PCA) of the 212 source rhesus macaques revealed strong separation between founding populations of animals (Fig. 2A). The conventional source animals are separated along PC1 (2.35% variance), and the Brooks and NEPRC animals are separated by PC2 (1.70% variance, Figure 2A). We performed an unsupervised admixture analysis on the 212 macaques using 2-5 theoretical ancestral populations (*K*) using ADMIXTURE v1.3.0 (Fig. 2B), with *K* = 3 best supported by cross-validation scores (SI Table S1). At *K* = 3, NEPRC animals are derived from a single ancestral population which is also observed at lower prevalence in both Brooks and conventional source animals. Both Brooks and conventional source animals also have a discrete component found at low frequency in the other population.

**Figure 2.**
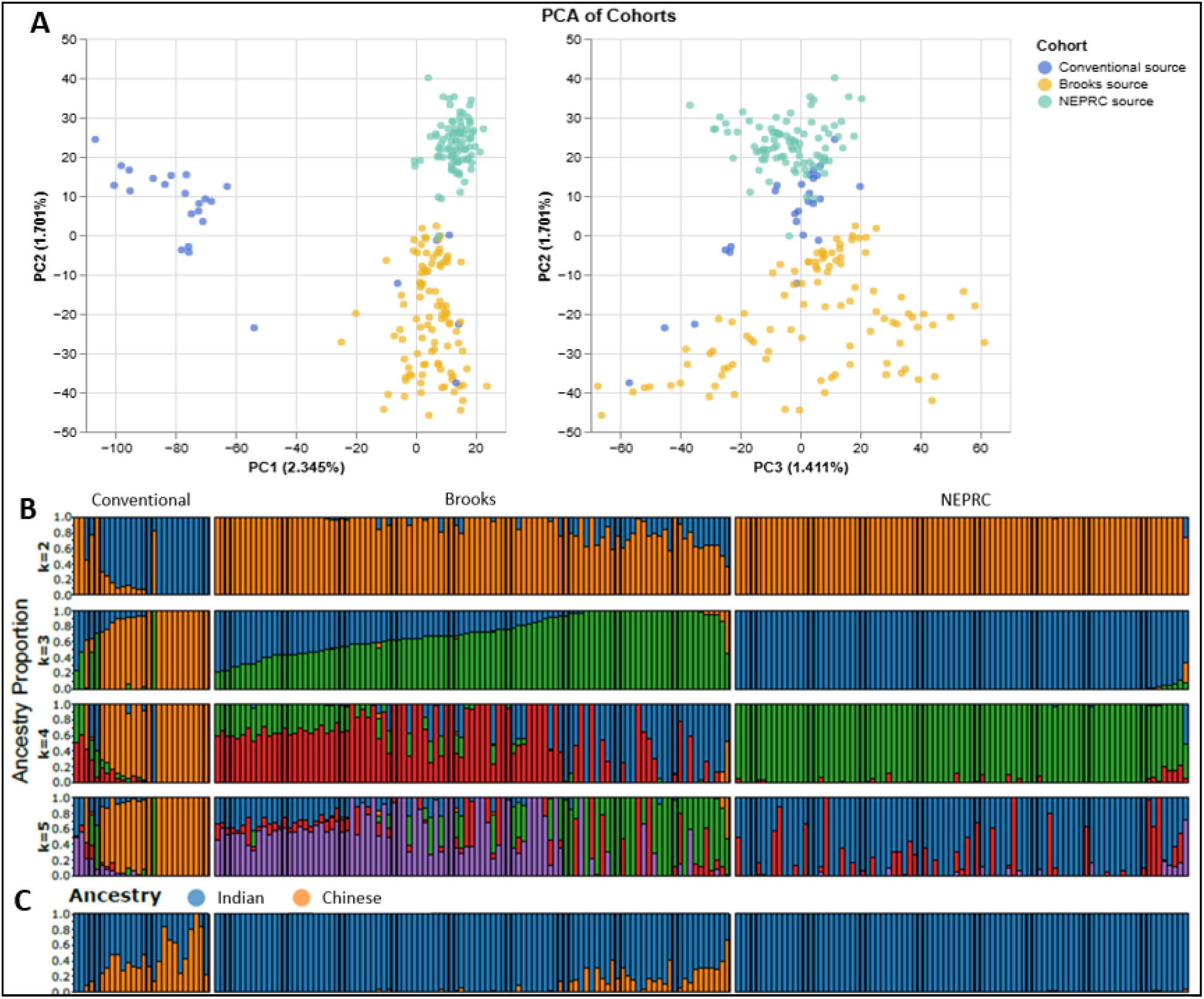
Admixture in SNPRC rhesus macaques. (A) Principal component analysis (PCA) of SNPRC rhesus macaques from three sources. (B) An unsupervised analysis of rhesus macaques (*n* = 212) grouped by source showing *K* = 2-5 theoretical ancestral populations. (C) A supervised analysis using macaque references genomes from animals of known Chinese- (*n* = 12) and Indian-origin (*n* = 12).

The genetic diversity of rhesus macaque populations in US-based biomedical research is driven by importation of animals from India and China. We performed a supervised ADMIXTURE analysis (Fig. 2) to investigate potential admixture between Chinese- and Indian-origin in the 212 source animals using reference sequence data for rhesus of known origin (Chinese *n* = 12, Indian *n* = 12) available through the macaque genotype and phenotype (mGAP) database (40). Among the conventional source animals, we identified one animal of Chinese-origin (99.9%) and three others that were highly admixed (>20% Chinese), with most of the remaining animals having some Chinese admixture (range 23-77%, SI Table S2). Among the Brooks source animals, which were purported to be non-admixed Indian origin, we identified 12 animals that were highly admixed (range 31-66% Chinese-origin) and 6 others with moderate levels of admixture (>15% Chinese-origin, Fig. 2C, SI Table S2). The NEPRC source animals showed negligible Chinese admixture (range 0.01-4%), suggesting the NEPRC founding animals were of Indian origin (Fig. 2C, SI Table S2). Notably, our admixture analysis is most strongly supported by *K*=3 ancestral populations suggesting that Chinese vs. Indian admixture is too simple a dichotomy to explain substructure.

We used WES data from the 212 source animals to calculate heterozygosity, fROH, and inbreeding coefficients to assess overall genetic diversity. The conventional source animals had significantly higher fROH and inbreeding than the Brooks animals (*t* (120) = 5.059, *p* = 1.533x10^-6^; *t* (123) = 3.271, *p* = 0.001) and NEPRC (*t* (109) = 5.023, *p* = 1.994x10^-6^; *t* (111) = 2.568, *p* = 0.01), suggesting lower overall genetic diversity. The average inbreeding coefficient was also significantly higher in the NEPRC animals than Brooks (*t* (184) = -2.647, *p* = 0.009) with no notable difference in heterozygosity or fROH.

Finally, we compared the empirical and genetic estimates of pairwise kinship for the source populations, as founding animals are presumed to be unrelated. We estimated pairwise relatedness for the 212 animals using KING v2.2.7 and uncovered several pairs of founding animals that were related. Notably, there were twelve pairs of Brooks source founders presumed unrelated that were actually first-order relatives (φ > 0.21, SI Table S3). There were also two conventional source males and two NEPRC animals that were first-order relatives (SI Table S3). All of these animals had offspring, resulting in animals of subsequent generations in the pedigree having higher kinship coefficients than estimated by the pedigree alone.

### Contemporary colony population genetics

#### Sequencing

To investigate the impact of management strategies aimed at increasing genetic diversity and decreasing admixture and the prevalence of certain MHC haplotypes in the colony over time, we further sequenced 276 rhesus macaques born into the SPF colony between 2003 and 2020 using WES. We divided these into three-year birth intervals for analyses to account for variation in numbers of animals sequenced per birth year (Table 1). We further divided animals born from 2018-2020 into two groups: one with two parents from the SPF SNPRC colony (birth cohort 6, *n* = 42) and one with one parent from NEPRC and one from the SNPRC colony (birth cohort 7, *n* = 41; Table 1) to assess potential genetic impacts of the merger. We acknowledge that birth cohort 5 has fewer animals than other cohorts which may have skewed results (Table 1).

**Table 1.** Group designations for analysis of rhesus macaque whole exome sequencing data.

| Group | Description | Year(s) | Sample size |
| --- | --- | --- | --- |
| Conventional source | Non-SPF SNPRC colony | < 2000 | 28 |
| Brooks source | Imported from BAFB | 2000-2002 | 97 |
| NEPRC source | Imported from NEPRC | 2015 | 87 |
| Birth cohort 1 | SNPRC colony born | 2003-2005 | 38 |
| Birth cohort 2 | SNPRC colony born | 2006-2008 | 36 |
| Birth cohort 3 | SNPRC colony born | 2009-2011 | 52 |
| Birth cohort 4 | SNPRC colony born | 2012-2014 | 53 |
| Birth cohort 5 | SNPRC colony born | 2015-2017 | 14 |
| Birth cohort 6 | SNPRC colony born | 2018-2020 | 42 |
| Birth cohort 7 | SNPRC colony born post-merger | 2018-2020 | 41 |

#### Genetic diversity

We calculated the heterozygosity, fROH, and inbreeding for the 276 rhesus macaques to analyze change over time. Results suggest genetic diversity and inbreeding have remained consistent over the past 20 years despite very little migration (Fig. 3). We also compared the three measures between the NEPRC merger founders and the merger offspring (birth cohort 7; 1 SNPRC parent, 1 NEPRC parent; Table 1) and report a significant reduction in fROH (*t* (126) = -2.184, *p* = 0.031) and inbreeding (*t* (126) = -3.587, *p* = 0.0005) in the merger offspring (Figure 3). We also report a significantly lower fROH (*t* (81) = 2.077, *p* = 0.041) in birth cohort 7 (Table 1) compared with contemporary birth cohort 6 (2 SNPRC parents; Table 1). These results suggest that the strategy to increase genetic diversity by merging the SNPRC and NEPRC groups was successful.

**Figure 3.**
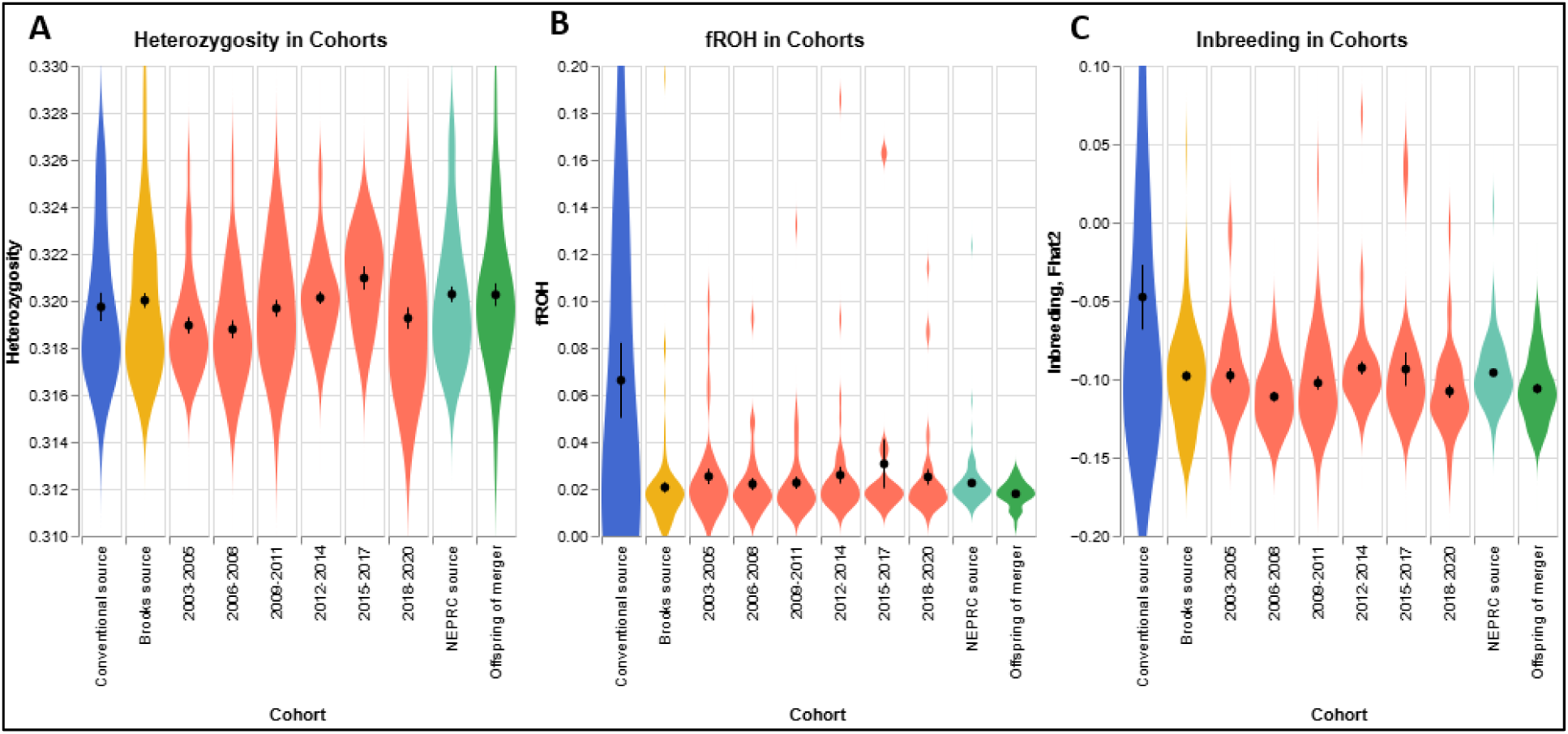
Genetic measures in the SNPRC rhesus macaques. (A) Heterozygosity measured as heterozygous SNV calls/called sites. (B) Fraction of the genome in runs of homozygosity (fROH). (C) Inbreeding based on excess homozygosity (F_hom_) of all founding and birth cohorts.

Breakdown of the groupings of SNPRC rhesus macaque whole exome sequencing (WES) data used for analyses. Birth cohort 7 had one parent each from SNPRC and NEPRC.

#### Admixture

To assess levels of Indian/Chinese admixture in the contemporary population, we ran a supervised analysis in ADMIXTURE on the 276 rhesus using the same reference animals as in the previous analysis (Chinese *n* = 12, Indian *n* = 12). Results indicate a gradual decrease over time in the number of animals above the admixed threshold of ≥15% Chinese ancestry, with very little admixture in the present-day population (SI Table S2, Fig. 4). Additionally, the average amount of Chinese ancestry in the admixed animals decreased from 30% in birth cohort 1 to 18% in birth cohort 7 (SI Table S2).

**Figure 4.**
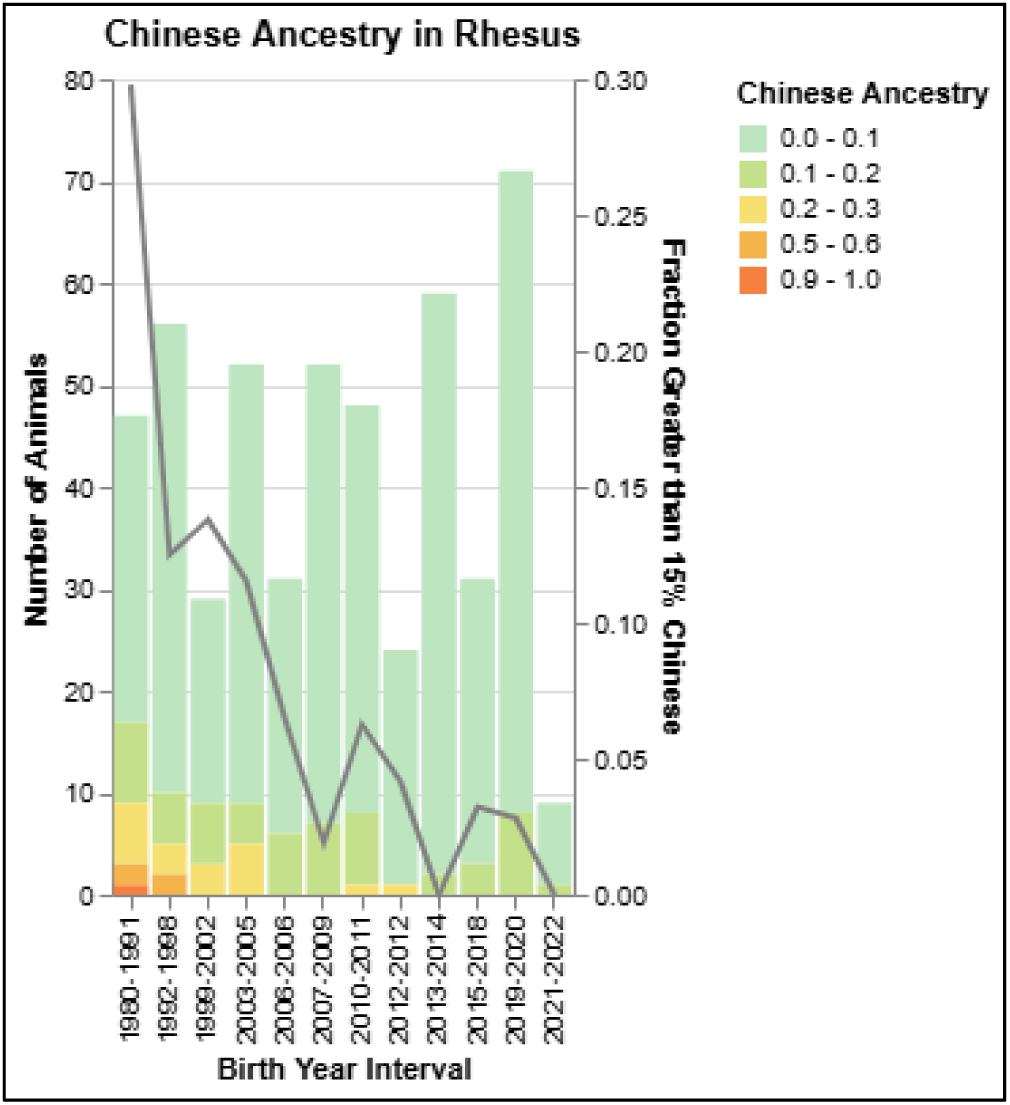
Proportion of Chinese admixture by birth year. Stacked bars show the percentage of Chinese ancestry by birth year interval and the grey line shows the fraction of the population with greater than 15% Chinese ancestry in the population over time.

#### Signatures of selection

The artificial selection placed upon the SNPRC animals to remove SIV-refractory MHC haplotypes is strong, with selection coefficients (*s*) of the reduction of specific haplotypes estimated to be -0.072, -0.088 and -0.078 for *Mamu-A001*, *Mamu-B008* and *Mamu-B017*, respectively. We explored temporal signatures of allele frequency variation across the pedigree to identify other regions of the genome which may be under selection in the SNPRC pedigree. We estimated F_ST_ in windows of 10,000bp across the genome, comparing each birth cohort to the founding Brooks population. Using a threshold of F_ST_>0.075 we identified 12 diverged windows in 2003-2005, which rose to 112 windows by 2018-2020 (Fig. 5A-D). When comparing Brooks animals to animals born between 2018 and 2020 we identified multiple regions of the genome where consecutive windows show evidence of divergence (Fig. 5E). Three of these windows are highlighted in Fig 5F-H. Notably, we can see that the divergence in each window increases over time, supporting that the signal is driven by a consistent pressure rather than the result of limited sampling. These F_ST_ peaks align with genes that have putative roles in modulating immune response (e.g., PRKAR2B, PAQR3, A1CF) or inflammatory pathways (e.g., PIK3CG, ANXA3, A1CF) in humans (Fig. 5F-H) (41–46).

**Figure 5.**
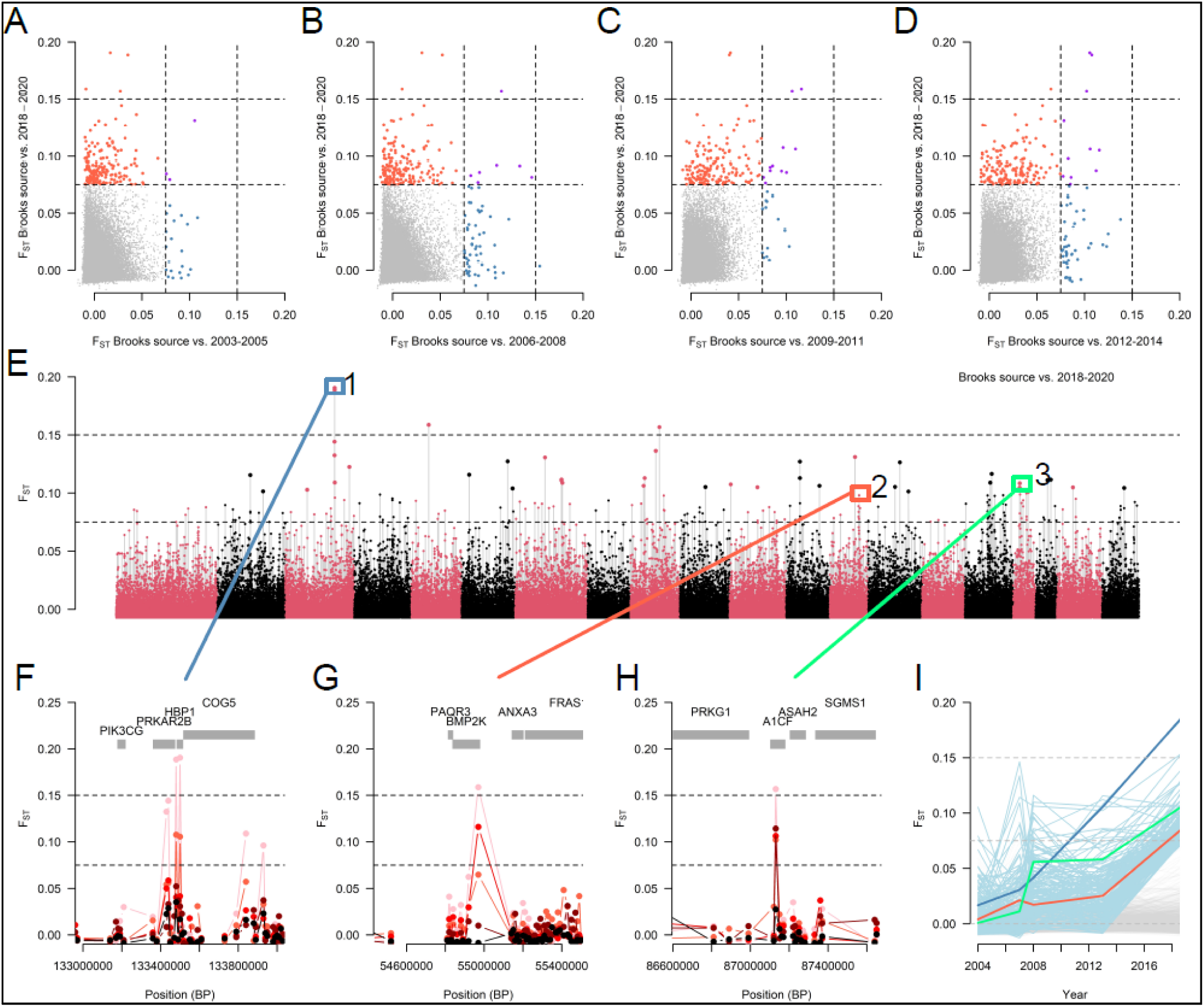
Short-term adaptation in the SNPRC macaque colony. We estimated FST in 10kb windows across exons comparing the founding Brooks colony animals to sequential time points. (A-D) The distributions of FST between latest and earlier time points. Windows with FST>0.075 are highlighted, red for Brooks vs. 2018-2020 only, blue for Brooks vs. additional time point, and purple for both. The intersection between timepoints increases over time. (E) Genomic distribution of FST between Brooks and the 2018-2020 birth cohort. (F-H) Three regions with multiple consecutive windows of FST>0.075. Each line shows FST between Brooks and 2003-2005 (black), 2006-2008 (dark red), 2009-2011 (red), 2012-2014 (light red), 2018-2020 (pink). (I) The trajectory of FST windows over time shows repeated signals at multiple loci (light blue lines). The regions in F-H are shown by matching colored lines. As 2015-2017 had few animals sampled they were not included in this analysis.

## Discussion

Here we have genetically characterized the primary rhesus macaque colony housed at SNPRC using MHC gene region and WES data. We estimated MHC haplotype diversity, genetic diversity, and admixture in the SNPRC rhesus macaques to assess the impact of various colony management strategies that have been implemented to ensure the continued supply of rhesus suitable for HIV/AIDS-related research. Overall, the colony has maintained relatively low levels of inbreeding and adequate levels of genetic diversity despite being closed to outside breeding for many years. We employ robust analyses of genetic and MHC diversity, relatedness, and inbreeding when making breeding recommendations. Previously these analyses were based primarily on pedigree information. However, recent advances in sequencing methodologies and decreased costs have allowed more accurate genomic characterization of NHP colonies designated for biomedical research.

Relating to MHC diversity, our results indicate that the removal or exclusion of animals with certain MHC haplotypes from breeding has not decreased the overall MHC haplotype diversity. Ongoing efforts to maintain MHC diversity include the inclusion of females with rare haplotypes in breeding, regardless of their status for *Mamu-A001*, *Mamu-B008*, and *Mamu-B017*, and plans to initiate cryopreservation of sperm from males with these and rare MHC haplotypes. While *Mamu-A001*, *Mamu-B008*, and *Mamu-B017* are undesirable for purposes of rapid SIV disease progression in HIV/AIDS research models (12, 13), the fact that they do confer some immune advantage is important in and of itself. Not enough is known about the complexities of the macaque MHC complex and how it impacts immune response more generally to completely eliminate these or any MHC haplotypes from captive rhesus populations. Future research into the specific mechanisms by which variation in MHC haplotypes leads to variable immune response will be critical for many biomedical research initiatives.

The Brooks source animals that founded the SPF breeding colony were purported to be of Indian-origin, however, our admixture analysis indicates that several animals had a substantial (> 15%) proportion of Chinese ancestry. Admixture of the offspring of un-sequenced Brooks source animals suggest that the number with Chinese ancestry was even higher. For example, rhesus with ID BS678 (Brooks source) sired ten animals in birth cohorts 2-7 (2006–2020) that showed greater than 15% Chinese ancestry (range 18-39%; SI Table S2), though no sample for DNA was available from this male. The dams of nine of the offspring were sequenced, and while one dam (CKDN220009637) showed 31% Chinese ancestry, the dams of the other offspring had negligible Chinese ancestry (86-99% Indian-origin; SI Table S2). These findings support other reports of cryptic admixture or even interspecific hybridization in early founding populations of captive NHP colonies (47–50). However, we still found relatively low levels of admixture overall in the contemporary rhesus population.

Beyond ancestral origin, our analyses revealed substantial population genetic substructure based on more recent animal source origins. Our PCA and admixture analyses suggest that demographic history and rapid shifts in allele frequencies led to distinct genetic signatures between animals in the three source populations. There were also differences in levels of genetic diversity between the source populations, with the conventional colony showing significantly lower levels of diversity than the Brooks and NEPRC source animals. The animals brought from the NEPRC also had significantly higher levels of inbreeding than the SNPRC animals. After merging these animals with the primary SNPRC breeding colony, we saw a significant decrease in fROH and inbreeding in the merger offspring compared to the NEPRC animals and the contemporary birth cohort with both parents from the same colony. This demonstrates that by merging isolated groups we can see significant improvement in genetic diversity within one generation. This has implications for increasing genetic diversity in both captive and wild animal populations.

We also investigated potential signatures of selection across the genome over time by comparing F_ST_ signals across birth cohorts with that of the Brooks source animals as they were the primary colony founders. Our F_ST_ analysis suggests there have been ongoing selective pressures driving changes in allele frequency over time. As we have been intentionally selecting against certain MHC haplotypes, and the MHC contains genes related to the adaptive immune system, we were not surprised to find F_ST_ peaks in several gene regions associated with immune response in humans. While none of these genes were located in the macaque MHC region, several play roles in supporting adaptive immune function, regulating immune response, modulating inflammation, or are expressed in immune related cells (41–46). Future work will further investigate signals of selection in these genes and their potential relationship to MHC-mediated immune function.

Here we demonstrate that WES is a useful methodology for monitoring genetic diversity and inbreeding in our rhesus macaque population while also providing invaluable exomic sequencing data for the investigation and potential development of macaque disease models of human health. The genetic characterization of NHPs designated for biomedical research has never been more accessible or more critical (51–56). It is not uncommon for NHPs of the same species from different geographical origins to show variation in disease susceptibility, progression and response (51). As noted above, rhesus macaque ancestral origins impact their suitability for HIV/AIDS-related research (28–30). Similar variation is seen in cynomolgus macaques (*Macaca fascicularis*), with animals from different geographical regions responding differently to *Shigella* immunization (57). Additionally, cynomolgus macaques historically hybridized with rhesus macaques in parts of their geographical range, and admixed cynomolgus susceptibility to malaria is strongly associated with their proportion of rhesus ancestry (50). These examples underscore the critical importance of genetically characterizing NHP populations designated for biomedical research and as translational models for human health.

### Conclusions

In this study we combined deep sequencing of the MHC region of ≈1,500 rhesus macaques with whole exome sequencing of ≈500 rhesus macaques for the genetic characterization of a captive colony designated for HIV/AIDS related research. These animals represent birth cohorts spanning more than 25 years, and include animals present at the founding of the SNPRC in 1999. We report that management strategies aimed at reducing the prevalence of SIV-refractory MHC haplotypes have been successful without reducing overall genetic or MHC diversity. We show that recent and historical demographic history has impacted population genetic structure, and we identify regions of the macaque genome with evidence of strong recent positive selection.

## Methods

### Study subjects

#### Historic SNPRC rhesus macaque breeding colony

The SNPRC was officially founded in 1999, though there were animals on site prior to that time comprising a conventional (non-SPF) colony of mixed Chinese and Indian ancestry. Between 2000 and 2002 the SNPRC received ≈250 presumed Indian origin animals from Brooks Air Force Base (BAFB) to establish the primary breeding colony. The two colonies were housed separately and the conventional colony was phased out over time, though some SPF derived animals were moved into the primary colony beginning in 2007. For the purpose of this study, we analyzed these data separately to represent the initial genetic makeup of the main colony (Brooks source) and as a supply of immigrant animals (conventional source).

#### NEPRC rhesus macaque colony merger

In 2015, the NEPRC disbanded and the SNPRC received 237 pedigreed, SPF Indian-origin rhesus that were colony-born at NEPRC. These animals were housed separately from the primary colony for approximately two years as they were acclimatized and space to merge the two groups became available. One of the primary reasons to merge the two groups for breeding was to increase overall genetic diversity, with the first offspring from mixed breeding groups born in 2018. At that time, the NEPRC group became part of the primary rhesus colony, which is now housed and managed as one. We generated WES data from ≈40% (*n* = 87) of the original NEPRC imports to estimate the baseline genetic diversity and admixture of the group.

#### Current rhesus macaque breeding colony

The SNPRC currently maintains a colony of ≈670 SPF animals that are designated primarily for HIV/AIDS-related research. Breeding animals are housed in one-male-multi-female groups or male-female pairs, so paternity is generally known. DNA samples are routinely extracted and banked for all animals that survive to weaning age (typically 9 months). The SNPRC has a large biorepository of DNA, blood, and tissue samples from most historical and all current NHP colonies, with the rhesus being the most current due to routine MHC haplotyping.

### MHC analyses

#### MHC sequencing

SNPRC started sporadic MHC genotyping in-house in 2007 using SSP-PCR to identify animals carrying *Mamu-A001*, *Mamu-B003*, *Mamu-B008*, and *Mamu-B017*. In 2013, we began sending samples to the WNPRC Genomics Services for MHC haplotyping by deep sequencing, with the 2012 birth cohort being the first with all individuals haplotyped. All NEPRC source animals were MHC haplotyped by WNPRC prior to arriving at SNPRC. In 2017 all animals with the *Mamu-B003* haplotype (*n* = 7) were removed from breeding and we began to limit the number of males with the other monitored haplotypes that were used to form new breeding groups. Females with monitored haplotypes have not been limited in order to maintain overall MHC diversity and rare haplotypes in the colony. Efforts to increase the production of animals that are considered “quad negatives”, being negative for all four monitored haplotypes, began in 2015, though efforts have increased recently with increased demand for animals for HIV/AIDS-related research.

#### MHC monitoring

We utilized existing MHC haplotype data for rhesus in the SPF colony (*n* = 1,458) to look at changes in haplotype diversity (*Mamu-A*, *Mamu-B*) and in frequency of the four haplotypes of interest (*Mamu-A001*, *Mamu-B003*, *Mamu-B008*, *Mamu-B017*) over time by birth cohort. Our analyses included animals born between 2012 (first full birth cohort fully typed) and 2022 (latest fully typed) inclusive. We measured haplotype diversity as 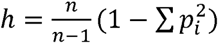 haplotypes with *p_i_* frequency. For the monitored haplotypes, we calculated the percent contribution of each of the four haplotypes to the total of all haplotypes present in the animals born during each year. We also calculated the percentages of total births that were quad negative for the same time period and compared across birth years.

### Genetic analyses

#### DNA sequencing

We extracted DNA from available blood and tissue samples using DNeasy Blood and Tissue Kits (Qiagen) following manufacturer’s instructions. DNA samples were then sent to Novogene (CA, USA) for QC, library prep, and whole exome sequencing. A fragment analyzer (Advanced Analytical Technologies, Inc.) was used for QC and library prep was completed using a human capture kit (Agilent SureSelect V6 60M Library Prep Kit). Final libraries were sequenced on the NovaSeq X Plus (Illumina) platform.

#### Data availability

Sequences have been deposited in NCBI under BioProject PRJNA810867. Additional data regarding the animals may be requested through a direct data intake request via the SNPRC website.

#### Alignment and variant calling

We completed alignment and genotype calling using existing bioinformatics pipelines. In brief, raw sequence data was trimmed using Cutadapt v4.0 (58) and aligned to the Mmul_10 rhesus macaque reference assembly using BWA-MEM v0.7.17 (59). After post-processing with SAMtools v1.21 (60) and base recalibration with GATK v4.2.3.0 (61), we called SNVs using GATK and joint-called with HaplotypeCaller v4.2.3 (62). SNVs were hard filtered to retain biallelic variants and filtered to exclude those matching the following: QUAL < 30.0, QD < 2.0, SOR < 3.0, FS < 5.0, MQ < 59.0, MQRankSum < -12.5, and ReadPosRankSum < -8.0. Genotype refinement (also with GATK) adjusted genotype scores and then filtered genotypes to exclude GQ < 30 and DP < 5. Finally, additional variant filtering removed sites with more than 20% of genotypes missing. Variants were annotated and consequences predicted using Ensembl Variant Effect Predictor v110 (63). Details of SNV results are available in SI Table S4.

#### Genetic diversity

We used WES sequencing data to analyze measures of genetic diversity in our contemporary breeding colony, as well as changes over time and before and after mergers. We divided the colony as follows: conventional source, Brooks source, NEPRC source, offspring born into the primary colony between 2003-2020 in three-year intervals, and offspring born after the NEPRC merger (one SNPRC and one NEPRC parent) between 2018-2020 (Table 1). We recognize that birth cohort 5 has far fewer samples than the others which could skew the results for that group. As a measure of overall genetic diversity, we calculated the heterozygosity and fractions of the genomes in runs of homozygosity (fROH) of each individual and averaged across each group using scikit-allel v0.20.3 (64). We also estimated the inbreeding coefficients for all animals based on excess homozygosity (F_hom_) using GTCA v1.94.1 (65).

#### Admixture analyses

We also used WES data to analyze genomic admixture in our colony in multiple ways. First, we assessed admixture between animals of Indian- and Chinese-origin across all groups described in Table 1 to identify original sources of Chinese admixture and to analyze change over time. Using the program ADMIXTURE v1.3.0 (66), we utilized genomic sequences of publicly available animals of known Indian- (*n* = 12) or Chinese-origin (*n* = 12) from mGAP (40) as references. Next, we investigated the impact of source (conventional, Brooks, NEPRC) on the genetic makeup of the population. We conducted a principal component analysis (PCA) and an unsupervised ADMIXTURE analysis using 2-5 theoretical ancestral populations (*K*).

#### Kinship estimation

The founding animals of a captive animal colony are those at the top of the pedigree that have no parentage information and are presumed unrelated. Births with known parentage are then used to create the rest of the pedigree, allowing kinship and relatedness to be analyzed for subsequent generations based on pedigree data alone. However, genetic data are required to determine potential relationships between the founding animals. Our Total Animal Care database houses all pedigree data and estimates pairwise kinship using the R tool kinship2. The kinship coefficient represents the genetic similarity of two individuals, with an expected value of 0.25 for first-order (e.g., parent offspring) relationships. We used WES data to estimate pairwise kinship between all sequenced animals using KING v2.2.7 (67), and compared those against the pedigree-estimated (empirical) kinship values.

#### Signatures of selection

We calculated the Weir & Cockerham F_ST_ using VCFtools and averaged across each birth cohort and source population in windows of 10,000bp across the genome, comparing each birth cohort to the founding Brooks population to investigate potential signatures of selection. Peaks with F_ST_>0.075 were considered regions of interest and we identified potential genes of interest in the most diverged windows, which corresponds to the 99.9^th^ percentile in the Brooks vs. 2018-2020 comparison. Birth cohort 5 (Table 1) was excluded from this analysis due to low sample size.

## Declarations

### Ethics approval

This research protocol was approved by the Texas Biomedical Research Institutional Animal Care and Use Committee #629MM.

### Author’s contributions

MML, SAC, IHC conceived and designed the study; SG, CC, LAC generated data; TC, SBG, CNR, DK coordinated rhesus macaque colony management and animal sampling; AB, DN, IHC, MML analyzed the data; MML, IHC wrote the manuscript. All authors read and approved the final manuscript.

## Acknowledgements

We would like to thank Thomas Butler, Laura Condel, David Elmore, Patricia Frost, Sharon Price, Eric Vallender, and John VandeBerg for providing valuable information relating to the history of the macaque colony. We would also like to thank Robert Lanford and Elda Mendoza for their contributions to the rhesus macaque colony.

